# Reducing resistance allele formation in CRISPR gene drives

**DOI:** 10.1101/150276

**Authors:** Jackson Champer, Jingxian Liu, Suh Yeon Oh, Riona Reeves, Anisha Luthra, Nathan Oakes, Andrew G. Clark, Philipp W. Messer

**Author notes:** Corresponding authors: JC, PWM. Equal Contribution (Jackson Champer, Jingxian Liu).

## Abstract

CRISPR gene drives can efficiently convert heterozygous cells with one copy of the drive allele into homozygotes, thereby enabling super-Mendelian inheritance. This mechanism could be used, for example, to rapidly disseminate a genetic payload through a population, promising novel strategies for the control of vector-borne diseases. However, all CRISPR gene drives tested have produced significant quantities of resistance alleles that cannot be converted to drive alleles and would likely prevent these drives from spreading in a natural population. In this study, we assessed three strategies for reducing resistance allele formation. First, we directly compared drives with the *nanos* and *vasa* promoters, which showed that the *vasa* drive produced high levels of resistance alleles in somatic cells. This was not observed in the *nanos* drive. Another strategy was the addition of a second gRNA to the drive, which both significantly increased the drive conversion efficiency and reduced the formation rate of resistance alleles. Finally, to minimize maternal carryover of Cas9, we assessed the performance of an autosomal drive acting in the male germline, and found no subsequent formation of resistance alleles in embryos. Our results mark a step toward developing effective gene drives capable of functioning in natural populations and provide several possible avenues for further reduction of resistance rates.

## INTRODUCTION

CRISPR gene drives have been proposed as a novel strategy for the control of vector-borne diseases by rapidly spreading alleles in a population through super-Mendelian inheritance^1-5^. Such a mechanism could be used, for example, to disseminate a genetic payload that reduces pathogen transmission in a disease vector population. Other potential applications include the suppression of agricultural pests or invasive species by spreading alleles that impair viability or fertility.

A CRISPR gene drive construct contains a Cas9 endonuclease that cleaves the genome at a target site specified by a guide RNA (gRNA). It then copies itself to that site via homology-directed repair (HDR). By this process, a heterozygote for the drive allele will be converted into a homozygote, enabling the rapid spread of such an in population^6-11^. The technical feasibility of CRISPR gene drive was first demonstrated in *Drosophila melanogaster*^12^ and *Saccharomyces cerevisiae*^13^. Subsequent studies have also applied this mechanism to the design of constructs aimed at population suppression in *Anopheles gambiae*^14^ and the spreading of a malaria resistance payload in *Anopheles stephensi*^15^.

The greatest unsolved obstacle to current CRISPR gene drive approaches is the formation of resistance alleles that cannot be converted to drive alleles^16,17^. Such resistance alleles can be produced by the drive itself when Cas9-induced cleavage is repaired by non-homologous end joining (NHEJ), instead of HDR, which will often change the sequence of the target site so that it is no longer recognized by the gRNA^12,14,15^. All homing drives tested in insects thus far have produced significant amounts of these resistance alleles. For example, in a previous study of a drive targeting the X-linked *yellow* gene in *D. melanogaster*, we observed that 29% of wild-type alleles were converted to resistance alleles in the germline of heterozygous females^18^. Furthermore, we found that resistance alleles frequently formed in the embryo due to persistence of maternally deposited Cas9. For instance, in daughters inheriting a wild-type allele from their father and a drive allele from their mother, we observed that 20% of the wild-type alleles were converted to resistance alleles. Such post-fertilization formation of resistance alleles occurred at an even higher rate in the *A. stephensi* study^15^. These high rates of resistance allele formation would almost certainly prevent a drive from spreading in a population, especially when it carries a fitness cost^17,19^.

In order to develop CRISPR gene drive as an effective means for genetic transformation of natural populations, the rates of resistance allele formation must be greatly reduced. One proposed strategy is to utilize constructs with multiple gRNAs targeting adjacent sites^3,4,20^. In this case, resistance would need to evolve at all sites to prevent drive conversion. In a two-gRNA drive, for example, if NHEJ-repair produces a resistance allele at the first site, Cas9 could then still cleave the second site, allowing insertion of the drive construct by HDR. Another strategy is to use a drive where conversion can occur in the male germline. Due to the small size of sperm, males would not transmit significant amounts of Cas9 into the embryo, thereby reducing postfertilization resistance allele formation. Finally, using a germline-restricted promoter to express Cas9 could reduce resistance allele formation in somatic cells. This could alleviate fitness costs in heterozygotes in systems targeting an essential/haploinsufficient gene, which has been proposed as an alternative strategy for resistance reduction^3,4,20^ and is a critical component of population suppression approaches.

In this study, we explore these three strategies by studying the performance of several CRISPR gene drive constructs in the model organism *D. melanogaster*. We first compare two constructs that target the X-linked *white* gene using the *nanos* and *vasa* promoters, demonstrating that leaky expression from *vasa* produces resistance alleles in somatic cells, which we do not observe in our drive using the *nanos* promoter. Second, we study a two-gRNA version of our *nanos* drive targeting *white*. The addition of a second gRNA significantly reduces formation of resistance alleles, particularly in the germline. Third, we study an autosomal drive targeting the autosomal *cinnabar* gene. This construct can perform drive conversion in the male germline with no subsequent formation of resistance alleles post-fertilization in the embryo.

## MATERIALS & METHODS

### Plasmid construction

The starting plasmids pCFD3-dU6:3gRNA^21^ (Addgene plasmid #49410) and pCFD4-U6:1_U6:3tandemgRNAs (Addgene plasmid # 49411) were obtained from Simon Bullock. Starting plasmids IHDyN1 and IHDyV1 were constructed in our previous study^18^. All plasmids were digested with restriction enzymes from New England Biolabs. PCR was conducted with Q5 Hot Start DNA Polymerase (New England Biolabs) according to the manufacturer’s protocol. Gibson assembly of plasmids was conducted with Assembly Master Mix (New England Biolabs) and plasmids were transformed into JM109 competent cells (Zymo Research). A list of plasmids and DNA oligo sequences used to construct them is detailed in the Supplementary Methods. Cas9 gRNA target sequences were identified using CRISPR Optimal Target Finder^22^.

### Generation of transgenic lines

The three gene drive lines in the study were transformed at GenetiVision by injecting the drive plasmid (BHDwN1, BHDwV1, BHDwN2, or BHDcN1) into individual Canton-S *D. melanogaster* lines. The drive plasmids were purified with Purify ZymoPure Midiprep kit (Zymo Research). To improve the efficiency of transformation, an additional source of Cas9 from plasmid pHsp70-Cas9^23^ (provided by Melissa Harrison & Kate O’Connor-Giles & Jill Wildonger, Addgene plasmid #45945), and an additional source of gRNA (BHDwg1, BHDwg2, or BHDcg1) was also included in the injection. Concentrations of gene drive/donor, Cas9, and gRNA plasmids were approximately 145, 37, and 38 ng/μL, respectively, in 10 mM Tris-HCl, 23 μM EDTA, pH 8.1 solution. To obtain a homozygous gene drive line, the injected embryos were first reared and crossed with wild type Canton-S flies. The progeny with dsRed fluorescent protein in the eyes, which usually indicated successful insertion of the gene drive, were selected and crossed with each other for two generations. The stock was considered homozygous at the drive locus when all male progeny were dsRed fluorescent for two consecutive generations.

### Fly rearing and phenotyping

Flies were reared at 25°C with a 14/10 hr day/night cycle. Bloomington Standard media was provided as food every 2-3 weeks. During phenotyping, files were anesthetized with CO2 and examined with a conventional stereo dissecting microscope. The white phenotype was declared when both eyes were fully white. Flies were considered “mosaic” if any discernable mixture of white and red color was observed in either eye. Fluorescent red phenotype, independent of the eye color phenotype, was observed using the NIGHTSEA system in the eyes, ocelli and abdomen. All experiments involving live gene drive flies were carried out using Arthropod Containment Level 2 (ACL-2) protocols at the Sarkaria Arthropod Research Laboratory at Cornell University, a quarantine facility certified by USDA APHIS for research up to ACL-3. Additional safety protocols regarding insect handling approved by the Institutional Biosafety Committee at Cornell University were strictly obeyed throughout the study, minimizing the risk of accidental release of transgenic flies.

### Genotyping

Flies were genotyped to obtain the DNA sequence at the *white* gRNA target site. Individual flies were first frozen and then ground in 30 μL of 10 mM Tris-HCl pH 8, 1mM EDTA, 25 mM NaCl, and 200 μg/mL recombinant proteinase K (Thermo Scientific). The homogenized mixture was incubated at 37°C for 30 min and then 95°C for 5 min. 1 μL of the supernatant was used as the template for PCR to amplify the *white* gRNA target site. DNA was further purified by gel extraction and sequenced with Sanger sequencing. ApE was used to analyze DNA sequence information (http://biologylabs.utah.edu/jorgensen/wayned/ape). Mosaic sequences were assessed with SangeranalyseR (http://github.com/roblanf/sangeranalyseR).

## RESULTS

### Construct design and generation of transgenic lines

We designed four CRISPR/Cas9-based gene drive constructs in *D. melanogaster* to explore different strategies for reducing resistance allele formation (Figure 1). Our first three constructs target the coding region of the X-linked *white* gene, the disruption of which causes a recessive and easily-identifiable white eye phenotype (w). The first construct uses Cas9 driven by the *nanos* promoter together with a single gRNA, driven by a U6:3 promoter (Figure 1A). The second construct has a similar design, except that the *vasa* promoter is used instead of *nanos* (Figure 1B). The third drive uses the *nanos* promoter, but with two gRNAs (one driven by a U6:3 promoter, the other by a U6:1 promoter), targeting two different sites (~100 nucleotides apart) in the coding sequence of the *white* gene (Figure 1C). The fourth drive targets the autosomal *cinnabar* gene instead of the X-linked *white* gene (Figure 1D). The disruption of *cinnabar* causes a recessive orange eye phenotype (cn). All four constructs also contain a dsRed gene driven by the 3xP3 promoter, which expresses a red fluorescent protein predominantly in the eyes and ocelli (although we also observed significant expression in the abdomen). This dsRed gene allows us to easily identify if a fly inherited the gene drive allele (D) by expressing the dominant red fluorescent phenotype.

**Figure 1.**
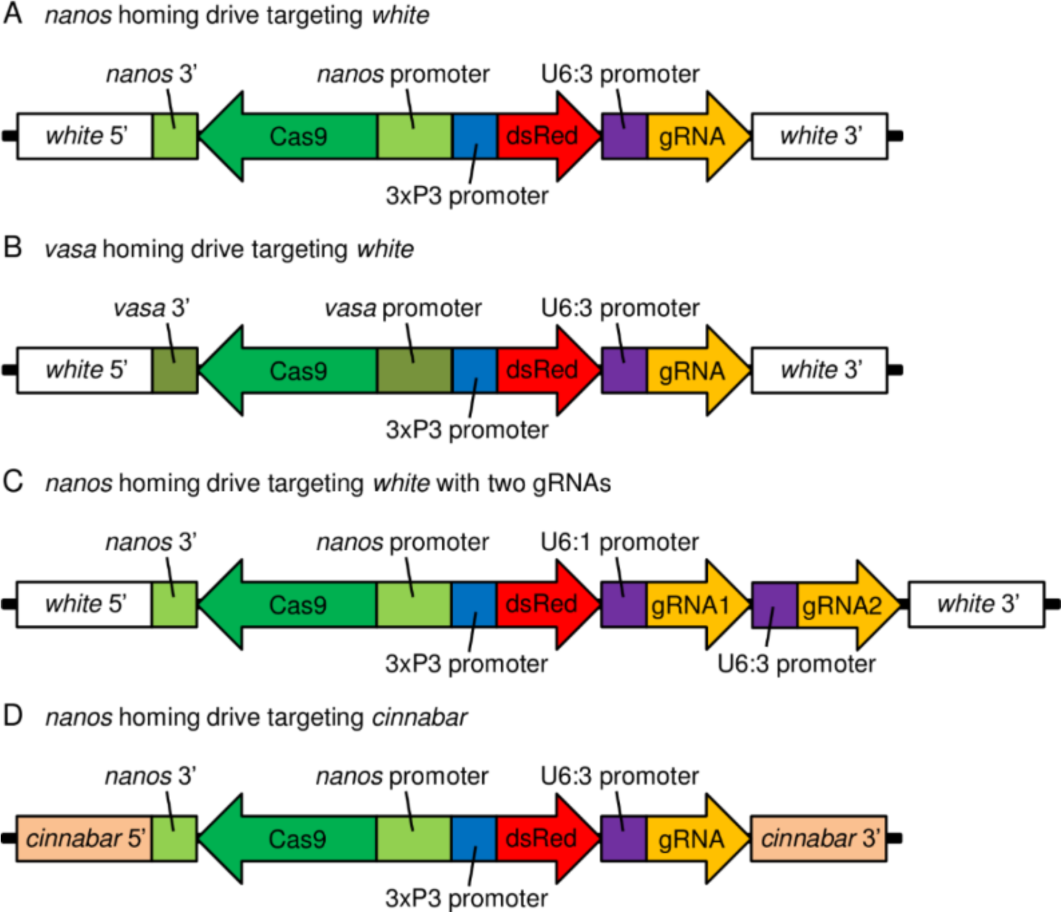
Gene drive constructs.

Since all four constructs target coding sequences, a successful insertion of any of the drives would disrupt the target gene. If the drive was not inserted and a resistance allele was formed, this should most likely result in a frameshift or sufficiently drastic change to the amino acid sequence, rendering the target gene nonfunctional (r2 resistance allele). Only occasionally would we expect a mutation at the target site to preserve the function of the target gene (r1 resistance allele). Box 1 summarizes the different phenotypes of our drive systems, together with the corresponding genotype combinations in male and female individuals.

##### Box 1. Genotypes, phenotypes, abbreviations.

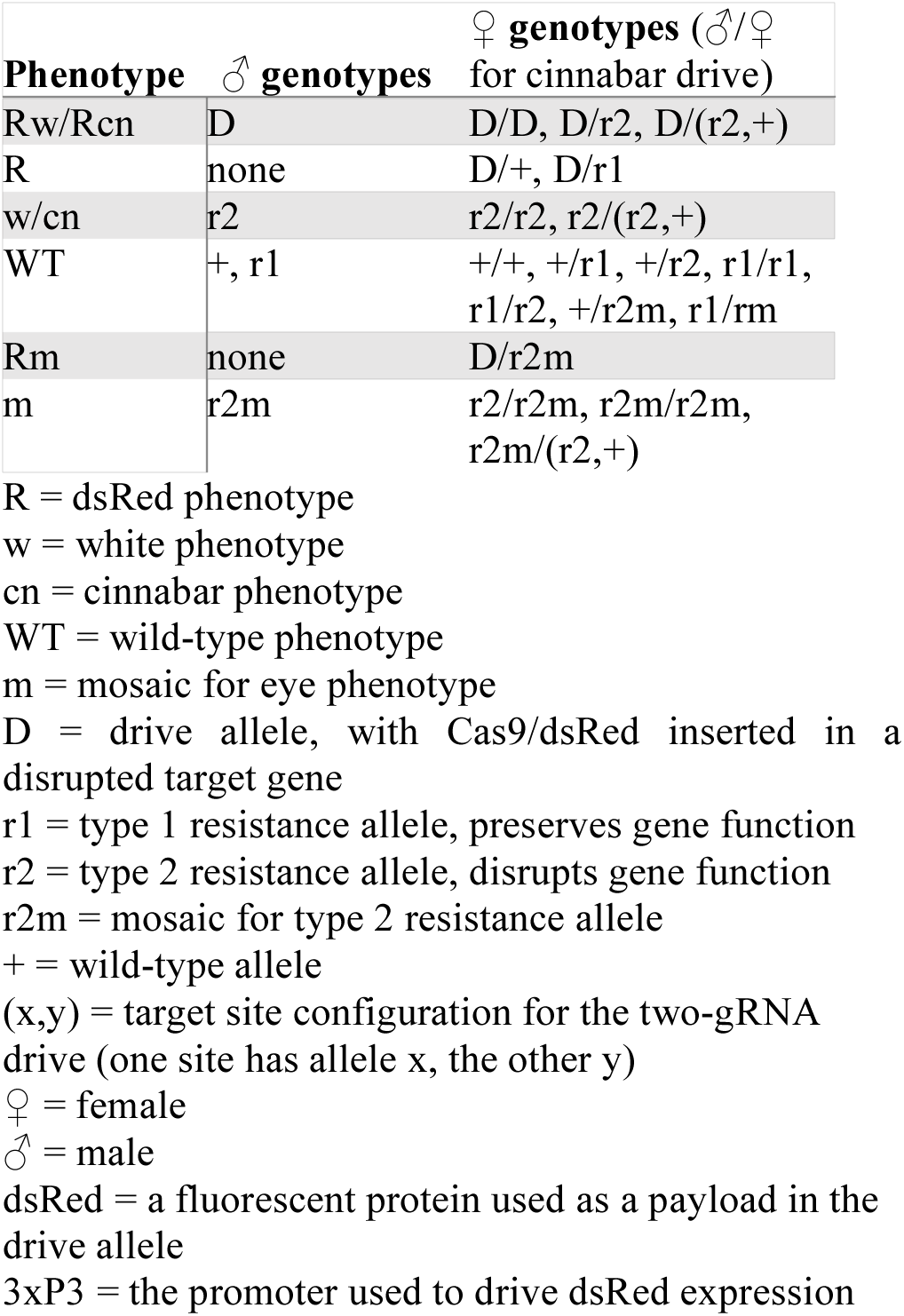

Isogenic fly lines were generated by injecting the gene drive plasmids into wild-type fly embryos from a Canton-S line. The flies were then reared, though no fluorescent phenotype was observed in the larvae or adults. All adults were then crossed with wild-type Canton-S flies, and several dsRed transformants with the gene drive were obtained. Most of these showed drive activity, but a small fraction did not, likely due to incomplete incorporation of the drive allele, and were excluded from the study.

### *Nanos* drive targeting *white*

Our *nanos* drive targeting the *white* gene appeared to follow a similar mechanism to our previous *nanos* drive targeting *yellow*^18^. When males with the drive allele were crossed with Canton-S females, the progeny followed Mendelian inheritance (Table 1A). No progeny exhibited the white phenotype, and only female progeny exhibited the dsRed phenotype (genotype D/+).

**Table 1.** Crosses of *nanos* drive targeting *white*.

A: Progeny of D males and +/+ females
|  | R | Rw | Rm | WT | w | m |
| --- | --- | --- | --- | --- | --- | --- |
| ♀ | 86 | 0 | 0 | 0 | 0 | 0 |
| ♂ | - | 0 | - | 78 | 0 | 0 |

B: Progeny of + males and D/+ females
|  | R | Rw | Rm | WT | w | m |
| --- | --- | --- | --- | --- | --- | --- |
| ♀ | 26 | 421 | 97 | 14 | 98 | 38 |
| ♂ | - | 650 | - | 7 | 146 | 2 |
| Total |  | 1194 |  |  | 305 |  |

C: Progeny of + males and D/D females
|  | R | Rw | Rm | WT | w | m |
| --- | --- | --- | --- | --- | --- | --- |
| ♀ | 5 | 122 | 16 | 0 | 0 | 0 |
| ♂ | - | 154 | - | 0 | 0 | 0 |
| Total |  | 297 |  |  | 0 |  |

D: Progeny of + males and D/r2 females
|  | R | Rw | Rm | WT | w | m |
| --- | --- | --- | --- | --- | --- | --- |
| ♀ | 13 | 70 | 18 | 4 | 52 | 11 |
| ♂ | - | 93 | - | 0 | 114 | 0 |
| Total |  | 194 |  |  | 181 |  |

To assess drive conversion efficiency, females with genotype D/+ were crossed to Canton-S males. dsRed phenotype was observed in 80% of the progeny, indicating that 59% of wild-type alleles were converted to drive alleles in the female germline (Table 1B). This is similar to the 62% conversion efficiency we observed for a similar drive targeting *yellow*^18^. In the same cross, 18% of male progeny had the white phenotype but not dsRed, indicating that 36% of wild-type alleles were converted to r2 resistance alleles in the germline, which was also similar to our previous study. Approximately 0.5% of male progeny had wild-type phenotype and inherited r1 resistance alleles. Sequencing of resistance alleles (Data S1) revealed that flies from the same mother were significantly more likely to have a sibling with an identical resistance alleles compared to an unrelated fly (*p*<0.0001, paired *t*-test). This suggests a mechanism in which resistance alleles are formed in germline stem cells, which can then give rise to multiple gametes sharing the same resistance allele.

In addition to the germline, resistance alleles can also form in the embryo after fertilization due to persistent maternal Cas9^24^. Such embryonic r2 resistance allele formation can result in a full white-eye phenotype or a mosaic eye color where r2 formation happened later in development in only a fraction of cells. Levels of conversion of paternal alleles to resistance alleles in the embryo post-fertilization were significantly higher in the *nanos* drive targeting *white* than in our previous drive targeting *yellow*^18^ (*p*<0.001, Fisher’s Exact Test). The level of embryonic r2 formation reached 77% in daughters inheriting the drive, as evidenced by daughters with phenotype Rw, with an additional 18% showing visible levels of mosaicism (Table 1B). Sequencing of six of these Rw females indicated that all had a single resistance sequence, in contrast to similar females from our *yellow* drive, which were each mosaic for different resistance allele sequences^18^. Among daughters that did not inherit a drive allele, only 68% were Rw, indicating formation of an r2 allele in the embryo after fertilization, though the number of mosaic flies (27%) was higher than for flies that did inherit a drive allele. Overall, this indicates that Cas9 expression was substantially higher for this drive than in our previous drive targeting *yellow*^18^, at least during the later expression phases, which produce Cas9 that persists into the embryo.

When females with genotype D/D were crossed to Canton-S males, the increased expression of Cas9 compared with D/+ females led to increased persistence of Cas9, and thus higher r2 allele formation rates in the embryo. Indeed, 85% of daughters of these flies were phenotype Rw, indicating conversion of paternal wild-type alleles into r2 resistance alleles (Table 1C), a significantly higher rate than in D/+ heterozygotes (*p*=0.038, Fisher’s Exact Test). Several Rw flies with phenotype Rw and genotype D/r2 were crossed to Canton-S flies, which resulted in approximately 50% of offspring inheriting a drive allele, as expected (Table 1D). Persistent Cas9, expressed at lower levels due to the presence of only one D allele, accounted for conversion of 69% of paternal alleles to r2 alleles in daughters, which was significantly lower than the rate in D/D females (*p*<0.0083, Fisher’s Exact Test). An additional 18% of daughters showed mosaicism.

### *Vasa* drive targeting *white*

Males with our *vasa* drive crossed to Canton-S females unexpectedly gave rise to daughters with full white or mosaic phenotype, in addition to dsRed phenotype (Table 2A). No phenotype R daughters were detected. Instead, all daughters had a white eye or mosaic phenotype. This contrasted with the *nanos* drive, which conformed to the Mendelian expectation that all daughters should have R phenotype. Such a result was also seen in a previous *D. melanogaster* gene drive targeting *yellow*^12^. A possible explanation is that leaky somatic expression of Cas9 cleaved the *white* gene in somatic eye cells, and NHEJ-repair resulted in a resistance allele with a disrupted white sequence and non-functional protein. If such leaky expression happened only in parts of the eye, then a mosaic phenotype would be observed.

**Table 2.** Crosses of *vasa* drive targeting *white*

| A: Progeny of D males and +/+ females |  |  |  |  |  |  |
| --- | --- | --- | --- | --- | --- | --- |
|  | R | Rw | Rm | WT | w | m |
| ♀ | 0 | 64 | 18 | 0 | 0 | 0 |
| ♂ | - | 0 | - | 67 | 0 | 0 |
| B: Progeny of + males and D/+ females |  |  |  |  |  |  |
|  | R | Rw | Rm | WT | w | m |
| ♀ | 2 | 222 | 21 | 1 | 43 | 27 |
| ♂ | - | 279 | - | 7 | 70 | 0 |
| Total |  | 524 |  |  | 148 |  |
| C: Progeny of + males and D/D females |  |  |  |  |  |  |
|  | R | Rw | Rm | WT | w | m |
| ♀ | 4 | 106 | 15 | 0 | 0 | 0 |
| ♂ | - | 134 | - | 0 | 0 | 0 |
| Total |  | 349 |  |  | 0 |  |
| D: Progeny of + males and D/r2 females |  |  |  |  |  |  |
|  | R | Rw | Rm | WT | w | m |
| ♀ | 0 | 100 | 15 | 0 | 53 | 30 |
| ♂ | - | 127 | - | 0 | 115 | 0 |
| Total |  | 242 |  |  | 198 |  |

However, all of these females appeared to remain D/+ in the germline, at least until normal germline Cas9 expression, which may happen concurrently with somatic expression. Drive conversion was shown to take place in the germline of these Rw females by crossing them to Canton-S males. dsRed phenotype was observed in 78% of progeny, indicating that the germline genotype remained D/+ until germline Cas9 expression, and that 56% of wild type alleles were then converted to drive alleles (Table 2B). This was approximately the level of the *nanos* drive and was slightly higher than the *vasa* drive targeting the *yellow* promoter from our previous study^18^. Phenotyping of male progeny indicated that 39% of wild-type alleles were converted to r2 alleles in the heterozygote female germline. Sequencing of resistance alleles (Data S2) showed the same pattern as the *nanos* drive, in which siblings were more likely to share a resistance allele sequence than unrelated flies (*p*=0.0021, paired *t*-test). Nearly all daughters of this cross that inherited a drive allele also showed the full white phenotype, which could be formed through either cleavage of the wild-type paternal chromosome by persistent maternal Cas9 in the early embryo or by later somatic expression of Cas9.

A similar pattern was seen when D/D homozygotes were crossed with Canton-S males, where almost all resulting daughters showed a full white phenotype (Table 2C). Maternally deposited Cas9 would be expected to form a D/r2 genotype early in the embryo, which would remain in the germline and prevent drive activity, while leaky somatic expression at a later stage should still allow normal levels of drive activity in the germline. To determine the germline genotype of these Rw daughters, twelve were crossed to Canton-S males. The progeny of nine of these showed only ~50% dsRed inheritance (Table 2D), indicating that the germline was mostly D/r2 (though some mosaicism may have been present). However, three Rw females showed high levels of drive activity, indicating their germline was initially D/+ (or mostly D/+) prior to the onset of germline Cas9 expression (Table S2B).

### Two-gRNA *nanos* drive targeting *white*

Our *nanos* drive with two gRNAs displayed qualitatively similar pattern as our *nanos* drive with one gRNA, but with improved drive efficiency. As with the one-gRNA drive, the crosses between gene drive males (D) and wild type females (+/+) followed Mendelian inheritance (Table 3A). When heterozygous females (D/+) were crossed with Canton-S males (+), dsRed phenotype was observed in 88% of the progeny, indicating that 76% of wild-type alleles were converted to drive alleles in the female germline (Table 3B). This is significantly higher than the one-gRNA *nanos* drive, where only 59% of wild-type alleles were converted to drive in D/+ females (*p*<0.001, Fisher’s Exact Test).

**Table 3.** Crosses of two-gRNA *nanos* drive.

|  |  |  |  |  |  |  |
| --- | --- | --- | --- | --- | --- | --- |
| **A: Progeny of D males and +/+ females** |  |  |  |  |  |  |
|  | R | Rw | Rm | WT | w | m |
| ♀ | 226 | 0 | 0 | 0 | 0 | 0 |
| ♂ | - | 0 | - | 249 | 0 | 0 |
| **B: Progeny of + males and D/(+,+) females** |  |  |  |  |  |  |
|  | R | Rw | Rm | WT | w | m |
| ♀ | 18 | 283 | 65 | 4 | 27 | 21 |
| ♂ | - | 467 | - | 0 | 60 | 0 |
| Total |  | 833 |  |  | 112 |  |
| **C: Progeny of + males and D/D females** |  |  |  |  |  |  |
|  | R | Rw | Rm | WT | w | m |
| ♀ | 0 | 176 | 8 | 0 | 0 | 0 |
| ♂ | - | 198 | - | 0 | 0 | 0 |
| Total |  | 382 |  |  | 0 |  |
| **D: Progeny of + males and D/(r2,r2) females** |  |  |  |  |  |  |
|  | R | Rw | Rm | WT | w | m |
| ♀ | 6 | 53 | 33 | 7 | 47 | 23 |
| ♂ | - | 124 | - | 0 | 93 | 0 |
| Total |  | 194 |  |  | 181 |  |
| **E: Progeny of + males and D/(r2,+) females** |  |  |  |  |  |  |
|  | R | Rw | Rm | WT | w | m |
| ♀ | 2 | 29 | 8 | 1 | 11 | 4 |
| ♂ | - | 50 | - | 0 | 22 | 0 |
| Total |  | 89 |  |  | 38 |  |

Since the two-gRNA drive had two target sites, if an r2 allele was formed at one site, drive conversion could still take place through cleavage at the second site. However, the drive conversion rate of 76% is significantly lower than the 83% predicted if the r2 alleles were formed independently at the same rate as in the one-gRNA drive, followed by conversion of any alleles with at least one wild-type site to D alleles (*p*=0.0045, Fisher’s Exact Test). This can potentially be explained by variable Cas9 expression levels (and thus cleavage rates) in individual cells, a reduction in HDR efficiency due to incompletely homologous ends around the remaining cleavage site, or a reduction in cleavage at individual target sites during the window for HDR due to the increased ratio of gRNAs to Cas9 in the germline cells. All of these factors may play a role in reducing the efficiency of multiple gRNAs.

The nature of the alleles formed in the maternal germline was assessed by phenotyping male progeny. None of the male progeny were wild type, indicating that all alleles had an r2 sequence for at least one of the two sites (Table 3B). This is a significantly lower rate than the one-gRNA drive (*p*=0.047, Fisher’s Exact Test), and can be accounted for by the fact that to create an r1 allele, repair at both sites would need to take place sequentially and would need to independently avoid disrupting the gene. A total of 11% of the male progeny had white phenotype with r2 alleles, indicating that 23% of wild-type alleles were converted to r2 alleles in the germline. Six of 26 alleles (23%) sequenced had an r2 allele at the first target site while the second had a wild-type allele (two of which subsequently underwent mosaic conversion to a resistance allele in the embryo). The asymmetry in r2 allele formation between the first and second target site could be due to the lower expression of the U6:1 promoter compared to the U6:3 promoter.

For the flies inheriting a drive allele, the rate of embryonic r2 resistance allele formation was 76%, with an additional 20% of flies showing mosaicism (Table 3B). The flies inheriting a wild-type allele had 50% embryonic r2 resistance allele formation, which was significantly lower (*p*=0.0003, Fisher’s Exact Test). An additional 43% of flies showed mosaicism. Overall, the embryonic r2 resistance rate of our two-gRNA construct was similar to our one-gRNA construct.

In crosses between Canton-S males (+) and homozygous drive females (D/D), 95% of the female progeny had white eye phenotype, and the rest displayed mosaicism (Table 3C). This was a significantly higher level of daughters with white phenotype than for D/+ mothers (*p*<0.0001, Fisher’s Exact Test). Most likely, the amount of Cas9 expressed in the eggs was increased due to increased gene copy number, so that more Cas9 persisted into the embryo, creating more resistance alleles.

To assess the genotype and drive characteristics of Rw females from the previous crosses, we crossed them with Canton-S males. Eight crosses showed little to no apparent drive conversion (Table 3D) and lower r2 allele formation in the embryo compared to D/D or D/+ females (*p*<0.0001, Fisher’s Exact Test). This indicated that their genotype was D/r2 (though there may have been some mosaicism, see below). However, the progeny from three crosses showed 70% inheritance for the gene drive, indicating a drive conversion rate of 40% (Table 3E). Drive conversion in Rw females was not observed in the one-gRNA *nanos* drive. In the two-gRNA drive, when a resistance allele formed at one target site and disrupted the function of the *white* gene, the other target site could potentially remain intact and allow conversion. Such a mechanism is supported by our sequencing results, which revealed that many of these flies had identical sequences at one cut site and variable sequences at the other (Data S3).

The conversion rate of 40% was significantly lower than the corresponding rate of 59% for the one-gRNA drive (*p*=0.017, Fisher’s Exact Test). It is possible that the imperfect match of homologous ends on one side of the target site reduces HDR repair efficiency. It is also possible that the second gRNA driven by the U6:1 promoter has lower activity than the first driven by the U6:3 promoter. Finally, the germline of these flies may actually have been mosaic for alleles that had r2 alleles at both target sites and alleles that still had a wild-type allele at one of the sites, reducing the apparent drive conversion rate.

### *Nanos* drive targeting *cinnabar*

In contrast to *white* and *yellow*, which are both X-linked, the *cinnabar* gene is autosomal. Our *nanos* construct targeting *cinnabar* should therefore be active in both the male and female germline. As with the other *nanos* drives, crosses between males homozygous for the drive allele (D/D) and Canton-S females (+/+) had progeny that followed Mendelian inheritance (Table 4A), all of which were genotype D/+.

**Table 4.**
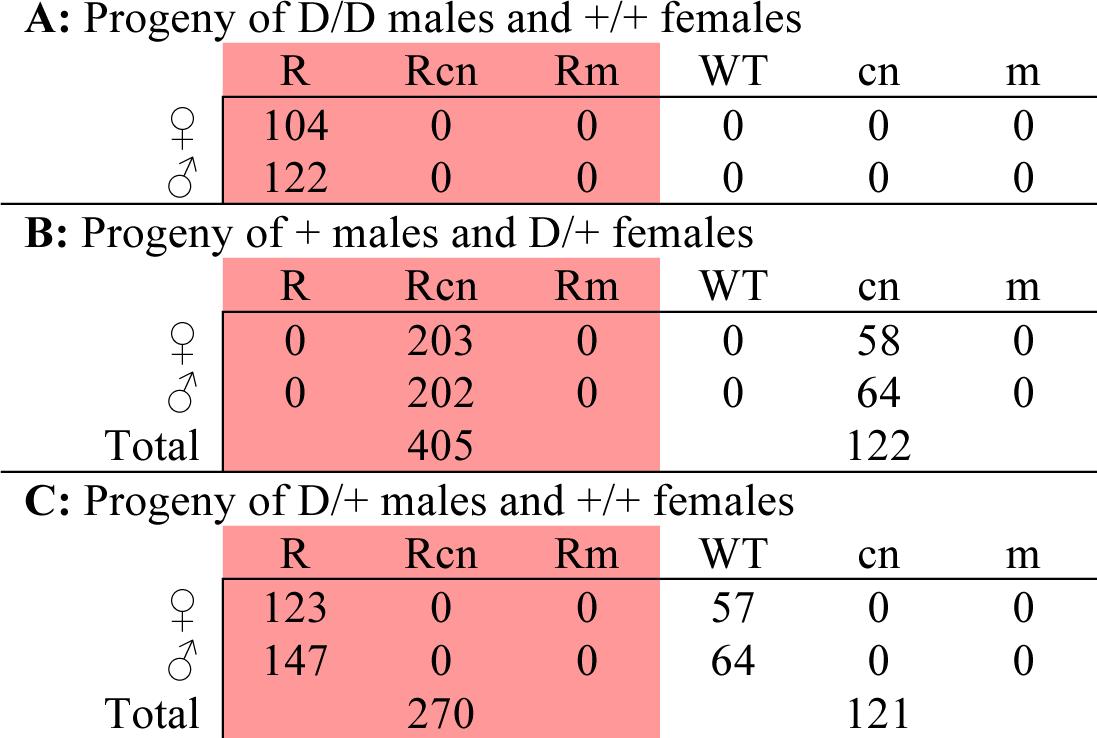
Crosses of *nanos* drive targeting *cinnabar*.

**A: Progeny of D/D males and +/+ females**
|  | R | Rcn | Rm | WT | cn | m |
| --- | --- | --- | --- | --- | --- | --- |
| ♀ | 104 | 0 | 0 | 0 | 0 | 0 |
| ♂ | 122 | 0 | 0 | 0 | 0 | 0 |

**B: Progeny of + males and D/+ females**
|  | R | Rcn | Rm | WT | cn | m |
| --- | --- | --- | --- | --- | --- | --- |
| ♀ | 0 | 203 | 0 | 0 | 58 | 0 |
| ♂ | 0 | 202 | 0 | 0 | 64 | 0 |
| Total |  | 405 |  |  | 122 |  |

**C: Progeny of D/+ males and +/+ females**
|  | R | Rcn | Rm | WT | cn | m |
| --- | --- | --- | --- | --- | --- | --- |
| ♀ | 123 | 0 | 0 | 57 | 0 | 0 |
| ♂ | 147 | 0 | 0 | 64 | 0 | 0 |
| Total |  | 270 |  |  | 121 |  |

Females with genotype D/+ from this cross were then mated to Canton-S males, and the progeny were assessed. Drive inheritance was 77%, as determined by dsRed fluorescence, for a drive conversion rate of 54%. This is similar to the conversion rate found in our one-gRNA *nanos* drive targeting *white*. Additionally, 100% of progeny from this cross had a cinnabar phenotype, indicating that almost all paternal chromosomes underwent cleavage by Cas9 and subsequent formation of r2 alleles. It is possible that a small fraction of the progeny was mosaic for the cinnabar phenotype, but this was difficult to visually assess. Taken together, these results imply that Cas9 expression in the *cinnabar* drive was substantially higher than for the *yellow* and *white* drives, both in the initial stages (at least in males) where resistance alleles may form in germline stem cells and especially later, resulting in higher levels of persistent Cas9 activity in the embryo after fertilization.

Males with genotype D/+ were crossed to Canton-S females, and drive conversion efficiency was found to be 38%, which was significantly lower than in females (*p*=0.01, Fisher’s Exact Test). No progeny had a cinnabar phenotype, indicating that a negligible amount of Cas9 persisted to the embryo, leading to reduced formation of resistance alleles in males by eliminating this second mechanism. However, sequencing of male wild-type progeny from this cross indicated that 62% of the wild-type paternal alleles had indeed been converted to resistance alleles in the male D/+ parental germline (Data S4).

### Modeling homing drive performance

Even though we were able to achieve lower resistance rates with our new drive constructs, it remains unclear whether these improvements are sufficient to enable a drive to reach high population frequency. To study this question, we modeled the frequency trajectories of drive, wild-type, and resistance alleles in panmictic populations under various drive scenarios, using our individual-based forward genetic simulation framework SLiM^25^. We assumed that drive alleles carried a (codominant) fitness cost of 5%, whereas resistance alleles had no fitness costs. Panmictic populations of 10,000 individuals were simulated, in which 100 homozygous gene drive flies were introduced at the start of each simulation. Allele frequency trajectories were then tracked over 40 discrete generations. We modeled both one-gRNA and two-gRNA approaches and autosomal versus X-linked drives. In each new offspring from a mother with a drive allele, every wild-type allele at any gRNA target site had a probability *e_1_* (if the mothers had one drive allele) or *e_2_* (if the mother had two drive alleles) of converting into a resistance allele. During the germline stage, a drive/wildtype heterozygote had a probability *g* of converting the wild-type allele into a resistance allele. If not converted, the wild-type allele could convert into a drive allele with probability *c*. The code for running these simulation analyses and accompanying data and figures are available at GitHub.

We studied three scenarios with different levels of Cas9 expression and resistance allele formation. (i) High (*e_1_*=0.75, *e_2_*=0.95, *g*=0.35, *c*=0.99 for one gRNA and *e_1_*=0.65, *e_2_*=0.8, *g*=0.3, *c*=0.95 for two gRNAs): designed to match our drives targeting *white*. (ii) Medium (*e_1_*=0.23, *e_2_*=0.32, *g*=0.3, *c*=0.98 for one gRNA and *e_1_*=0.2, *e_2_*=0.27, *g*=0.28, *c*=0.94 for two gRNAs): designed to match our drives targeting *yellow* from our previous study^18^. (iii) Low (*e_1_*=0.03, *e_2_*=0.05, *g*=0.25, *c*=0.97 for one gRNA and *e_1_*=0.03, *e_2_*=0.05, *g*=0.25, *c*=0.93 for two gRNAs): inspired by a GDL line from our earlier study^18^ that assumes a lower level of Cas9 expression and subsequently significantly reduced resistance allele formation in the embryo. All scenarios are presented in Figure 2.

**Figure 2.**
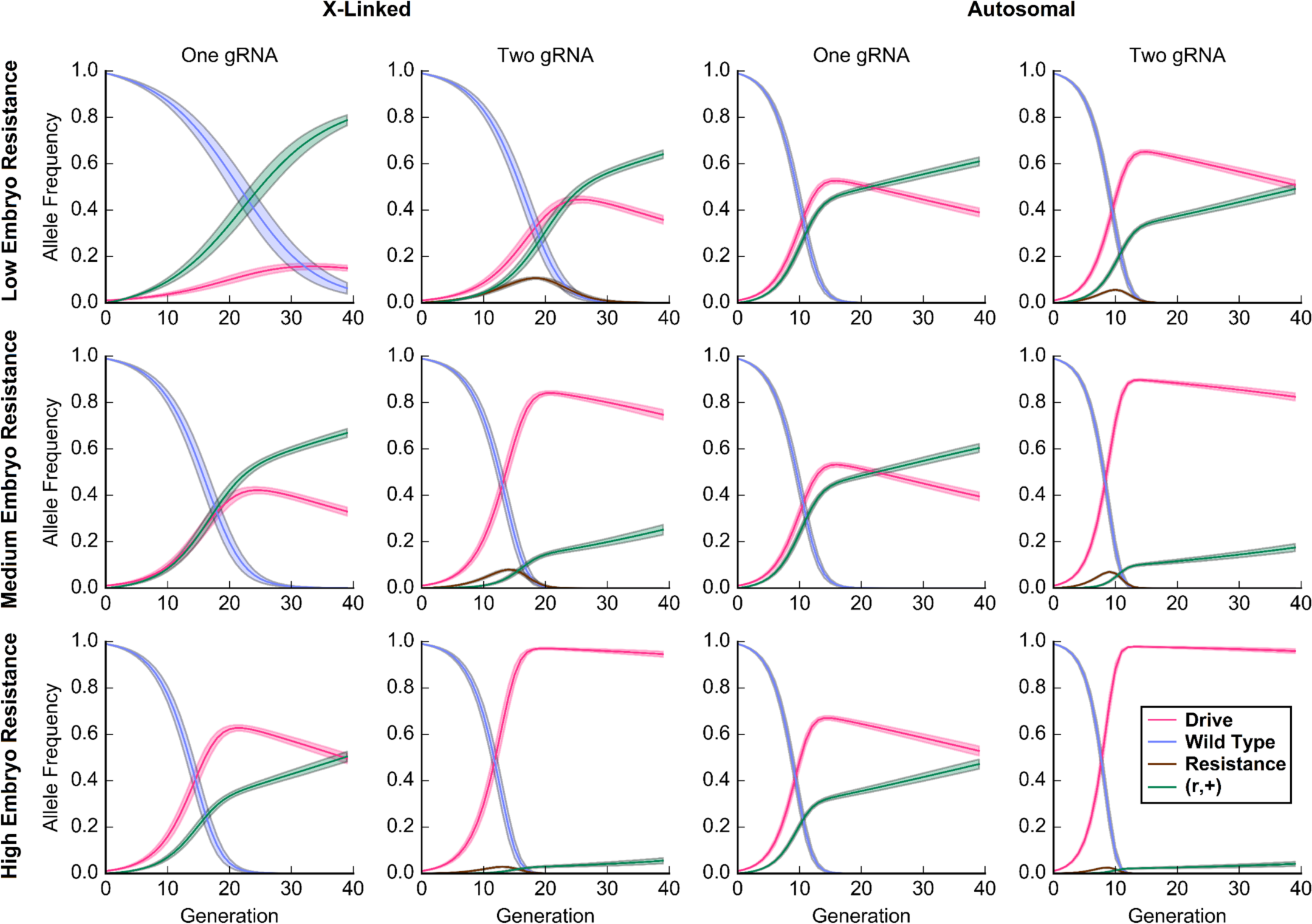
Simulated allele frequency trajectories of different drive systems. The panels show various drive models with different levels of resistance allele formation. The high-level scenario is inspired by our *white* drives, the medium level scenario by the *yellow* drives from our previous study, and the low-level scenario specifies a hypothetical drive with reduced Cas9 expression and persistence. The curves show the mean (solid lines) and standard deviation (shaded regions) of allele frequency trajectories averaged across 1,000 trials.

An X-linked drive similar to our *yellow* drive (medium resistance scenario) would only reach a maximum frequency of 42% after 24 generations in our idealized panmictic population model, before declining due to its fitness disadvantage compared to resistance alleles. A drive with the characteristics of our system targeting *white* (high resistance scenario) would only reach a maximum frequency of 16% after 33 generations. The addition of a second gRNA to the *white* system substantially improves the maximum drive allele frequency to 45% after 26 generations. A hypothetical, autosomal two-gRNA drive with a lower level of Cas9 expression (low resistance scenario) would reach a maximum frequency of 98% after 13 generations, falling to 96% after 40 generations.

These models show that to achieve a >90% drive allele frequency, even in the idealized scenario of a panmictic population we simulated here, at least two gRNAs are necessary, and resistance allele formation in the embryo must be further reduced from the levels we observed in our drives. Reducing germline resistance rates would also be beneficial, but we did not consider significant reductions at this stage since it would require a different promoter than either *nanos* or *vasa*, or a more complicated drive system.

## DISCUSSION

In this study, we explored three different strategies for reducing the rate of resistance allele formation for CRISPR gene drives. First, we tested differences in resistance mechanisms between constructs using the *nanos* and *vasa* promoters for Cas9 expression. Second, we assessed whether the use of multiple gRNAs targeting different sites can improve drive efficiency and reduce resistance allele formation. Finally, we assessed the performance of a drive targeting an autosomal site, in which drive conversion can also occur in males.

Our study builds upon a previous study in which we demonstrated drive conversion takes place in the germline with the *nanos* and *vasa* promoters, while resistance alleles can form in both the germline and postfertilization in the embryo by persistent maternal Cas9^18^. The present study allowed us to further refine these mechanisms (Figure 3). Specifically, we showed that resistance alleles of different offspring derived from the same parent are often identical, suggesting that resistance alleles can form in germline stem cells prior to meiosis and HDR, which then gives rise to multiple gametes. Table 5 summarizes the drive conversion and resistance allele formation rates we measured for the various drives studied here and in our previous study^18^.

**Figure 3.**
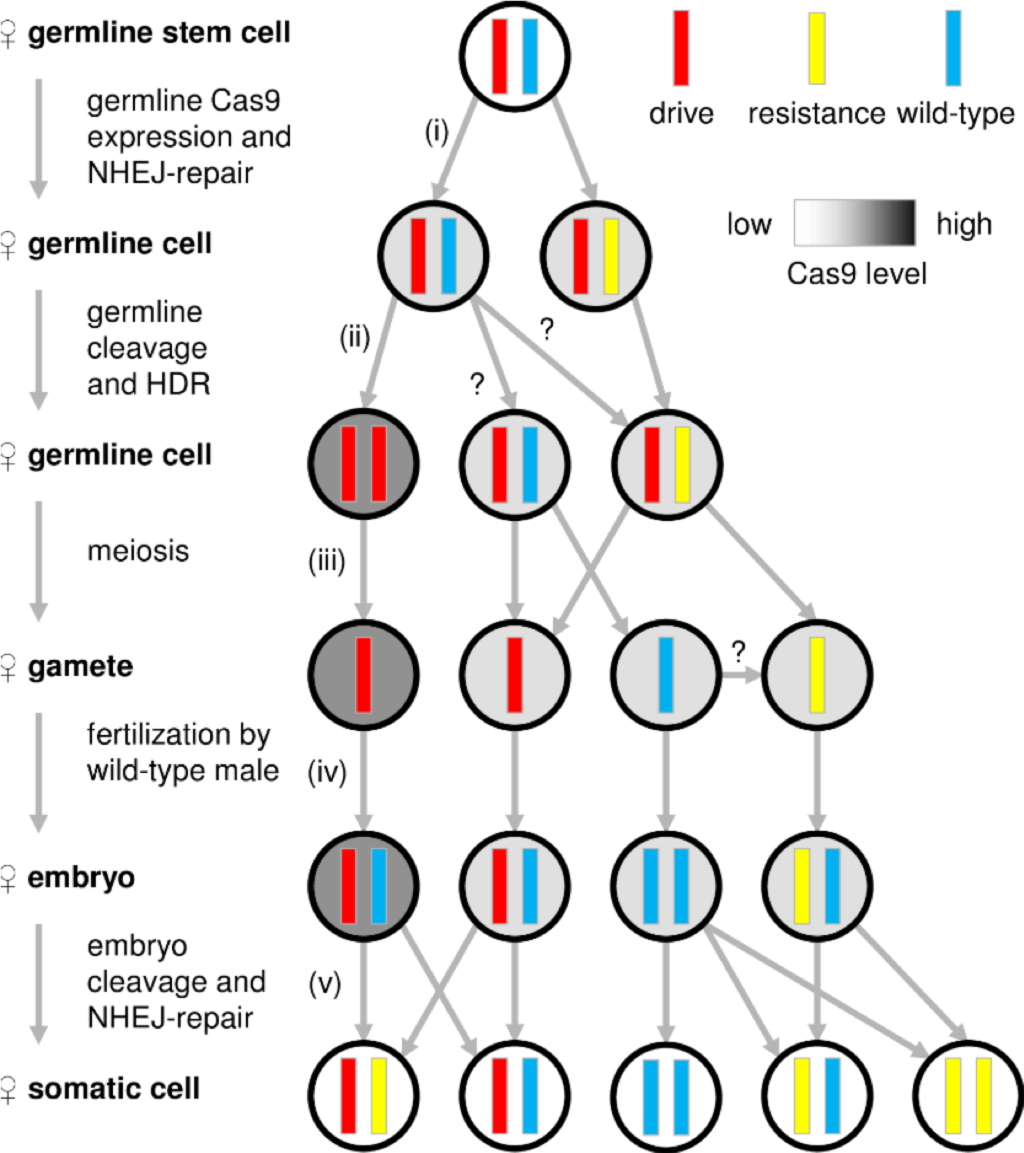
Mechanisms of resistance allele formation. (i) In a heterozygous female with genotype D/+, early expression of Cas9 in germline stem cells prior to the window for HDR can convert a fraction of wild-type alleles to resistance alleles by NHEJ. (ii) Nearly all remaining wild-type alleles will be converted to drive alleles by HDR prior to meiosis. (iii) Meiosis than takes place, and (iv) gametes undergo fertilization, in this example by a wild-type male. (v) After fertilization, persistent Cas9 can convert the paternal chromosome and any remaining maternal wild-type chromosomes to resistance alleles by NHEJ. We did not observe any successful drive conversion during this stage.

**Table 5.** Drive parameters for several *D. melanogaster* gene drives and fly lines.

| Fly line | Cas9 promoter | Target gene | Conversion rate of drive | Germline r2 formation rate | Embryo r2 formation rate |
| --- | --- | --- | --- | --- | --- |
| Canton-S | *nanos* | *white* | 59% | 36% | 77% |
| Canton-S | *vasa* | *white* | 56% | 39% | ~75% |
| Canton-S | *nanos* | *white-2gRNA* | 76% | 23% | 76% |
| Canton-S | *nanos* | *cinnabar* | 54% | 46% | ~100% |
| Canton-S | *nanos* | *cinnabar* (male) | 38% | 62% | 0% |
| *w<sup>1118</sup>* | *nanos* | *yellow* | 62% | 29% | 20% |
| Canton-S/*w<sup>1118</sup>* | *nanos* | *yellow* | 55% | 35% | 19% |
| GDL-Z/*w<sup>1118</sup>* | *nanos* | *yellow* | 54% | 39% | 28% |
| GDL-T/*w<sup>1118</sup>* | *nanos* | *yellow* | 51% | 41% | 56% |
| GDL-B/*w<sup>1118</sup>* | *nanos* | *yellow* | 50% | 36% | 26% |
| GDL-N/*w<sup>1118</sup>* | *nanos* | *yellow* | 46% | 52% | 22% |
| GDL-I/*w<sup>1118</sup>* | *nanos* | *yellow* | 40% | 47% | 4.2% |
| *w<sup>1118</sup>* | *vasa* | *yellow* Promoter | 53% | 47% (r1) | ~20% |
| Canton-S/*w<sup>1118</sup>* | *vasa* | *yellow* Promoter | 37% | 63% (r1) | ~20% |

r1 = an estimate for type r1 alleles are displayed for the drive targeting the *yellow* promoter, since r2 alleles are very rarely created at this site. Parameters for the drives targeting *yellow* are from our previous study<sup>18</sup>.

We further showed that the *vasa* drive frequently induces leaky somatic expression (Figure 4). This mechanism would provide an explanation for the apparent contradiction between our finding that *vasa* drives in the germline and a previous study of a similar *vasa* drive targeting *yellow*, which was thought to drive in the embryo^12^. That conclusion was based on the observation that the recessive yellow phenotype was observed in offspring that inherited one drive and one wild-type allele from their parents^12^. However, leaky somatic expression of *vasa* can equally produce this outcome, even in females that remain D/+ heterozygotes in the germline. A recent study in *A. stephensi* also showed some evidence of somatic expression from the *vasa* promoter, although at a much lower rate than we observed in *D. melanogaster*^15^.

**Figure 4.**
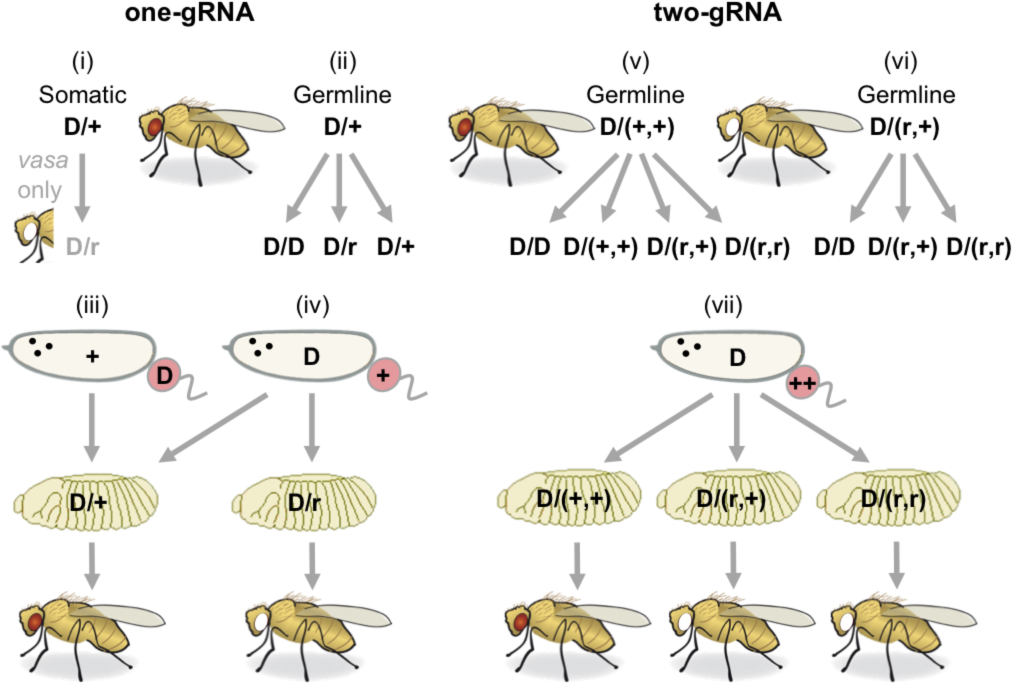
Comparison of resistance mechanisms between one-gRNA and two-gRNA drives. (i) In a D/+ female, leaky somatic expression in *vasa* drives can convert wild-type (+) alleles to resistance (r) alleles in the body, resulting in white eye phenotype, though drive conversion can still occur successfully in the germline. Such somatic expression did not occur in our *nanos* drives. (ii) Germline Cas9 expression can lead to successful drive conversion, formation of an r allele, or persistence of the + allele (unlikely). In males, no conversion can occur in the germline for our X-linked drives, but can occur for the autosomal *cinnabar* drive. (iii) Embryos derived from paternal D gametes will not experience maternal Cas9 cleavage. (iv) However, any gamete derived from a female with a D allele may contain sufficient Cas9 to convert a paternal + allele into an r allele. (v) A female with the two-gRNA drive with genotype D/(+,+) could convert the (+,+) allele into a D, an (r,+), or an (r,r) allele. (vi) In some cases, a fly with the two-gRNA drive will have a white phenotype and genotype D/(r,+), but still be able to utilize the remaining wild-type target site for successful homing. (vii) Maternal Cas9 can convert a paternal (+,+) allele to (r,+) or (r,r) alleles in the embryo.

Our *nanos* and *vasa* drives targeting the *white* gene had very similar conversion and resistance rates (Table 5). This contrasts with our previous study, in which the *nanos* drive offered a modest improvement in the drive conversion rate^18^. However, despite similar performance, leaky somatic expressions of the *vasa* promoter should still render *nanos* a superior choice for many applications, such as the targeting of haploinsufficient or fertility genes, where somatic activity could induce strong fitness costs.

The use of multiple gRNAs constitutes a promising strategy for improving the efficiency of gene drives and reducing resistance rates. In our study, we found that the addition of a second gRNA considerably increased the drive conversion efficiency of our *white* construct (Table 5). However, this improvement remained lower than would be expected in a model where resistance alleles form at each target site independently, and drive conversion at each individual site occurs at the same rate as observed in the one-gRNA construct. One possible explanation for this discrepancy is that additional gRNAs saturate Cas9, resulting in lower cleavage at each individual site, as indicated by the persistence of a higher number of wild-type sequences in the two-gRNA drive (Data S3) compared to the one-gRNA drive (Data S1). It is also possible that increasing distance between target sites could reduce drive efficiency (in our drive, the two target sites were ~100 nucleotides apart). In many genes, it would probably be possible to find up to four gRNA target sites with low expected off-target effects within a relatively small 200 nucleotide window. Methods to efficiently multiplex this number of gRNAs already exist^26-28^. This would potentially allow for further improvement of drive efficiency.

Importantly, we observed many instances of simultaneous cuts at both target sites in our two-gRNA drive, which led to the deletion of the region between the sites after NHEJ. In drives with more than two gRNAs, this mechanism could thus remove interior target sites, resulting in much higher resistance rates compared to an idealized system where no simultaneous cuts occur. Additionally, in genetically diverse wild populations such as *A. gambiae*^29^, finding several gRNA target sites with low off-target effects within a small window would be considerably more challenging. One possibility to address this problem would be to use higher fidelity versions of Cas9 with significantly lower off-target effects^30^, allowing targeting of sequences similar to other locations in the genome with lower fitness cost to the organism.

The use of multiple gRNAs may function particularly well with a strategy of targeting a haploinsufficient gene and reforming the gene as part of successful HDR^3,4,20^. In such a system, embryos with r2 alleles would be non-viable. The use of multiple gRNAs would reduce the chance of r1 alleles being formed, since the target sites would each need to acquire an r1 sequence independently. Additionally, these r1 sequences may be low in diversity, allowing them to be targeted by additional gRNAs incorporated into the original gene drive or in a follow-up drive. However, a multiple-gRNA system targeting a haploinsufficient gene may still need to address the issue of incomplete HDR repairing the target gene region but not inserting the drive, thus forming an r1 allele. While this occurs at a low rate, we did observe several instances of small insertions from incomplete HDR (Data S3, Data S4, previous study^18^), and our genotyping method was not optimized to detect larger insertions.

An alternative (or complementary) strategy to multiple-gRNA drives for reducing the formation of resistance alleles would be an improved promoter. The *nanos* and *vasa* promoters in *D. melanogaster* begin expression of Cas9 earlier than the window for HDR, resulting in undesirable formation of resistance alleles in germline stem cells. The *vasa* promoter in the *A. stephensi* gene drive system^15^, on the other hand, appears to largely avoid this problem. It is possible that the *nanos* promoter would function similarly well in this species. However, all of these systems still have high levels of maternal Cas9 persistence through the embryo stage, resulting in high levels of additional resistance allele formation. An ideal promoter would express Cas9 only during the window for HDR, which would then be degraded rapidly prior to fertilization. A male-only promoter would avoid the issue of maternally deposited Cas9 as shown by our *cinnabar* drive, which formed no resistance alleles in the offspring of males with the drive. However, such a promoter would still need to avoid expression of Cas9 before the window for HDR. These issues could also be mitigated if the gRNAs were expressed in the germline only during the window for HDR, in lieu of the ubiquitous promoters that have been used in all CRISPR gene drives thus far. Indeed such multiplexing systems capable of being driven by a variety of promoters have already been developed using ribozyme-based methods^26,28^. However, a significant increase in the size of the drive constructs from such promoters may prove to be a disadvantage, particularly if payload genes are also very large.

Our study emphasizes that resistance will likely remain the prime challenge facing CRISPR gene drives, but also demonstrates that a two-gRNA approach can substantially reduce resistance allele formation. While the use of multiple gRNAs by itself will likely be insufficient to create drives that are efficient enough for use in wild populations, they will probably be a critical part of a successful strategy.

## ACKNOWLEDGEMENTS

We thank Sam Champer for assistance with statistical analysis and Chen Liu for assistance with sequencing resistance alleles. This research was supported by startup funds from the College of Agriculture and Life Sciences at Cornell University to P.W.M and the Meinig Family Investigator award to A.G.C.

## SUPPLEMENTARY INFORMATION

### Supplementary methods

The following tables show the DNA fragments used for Gibson Assembly of the listed plasmids. PCR products are shown with the oligonucleotide primer pair used, and plasmid digests are shown with the restriction enzymes used.

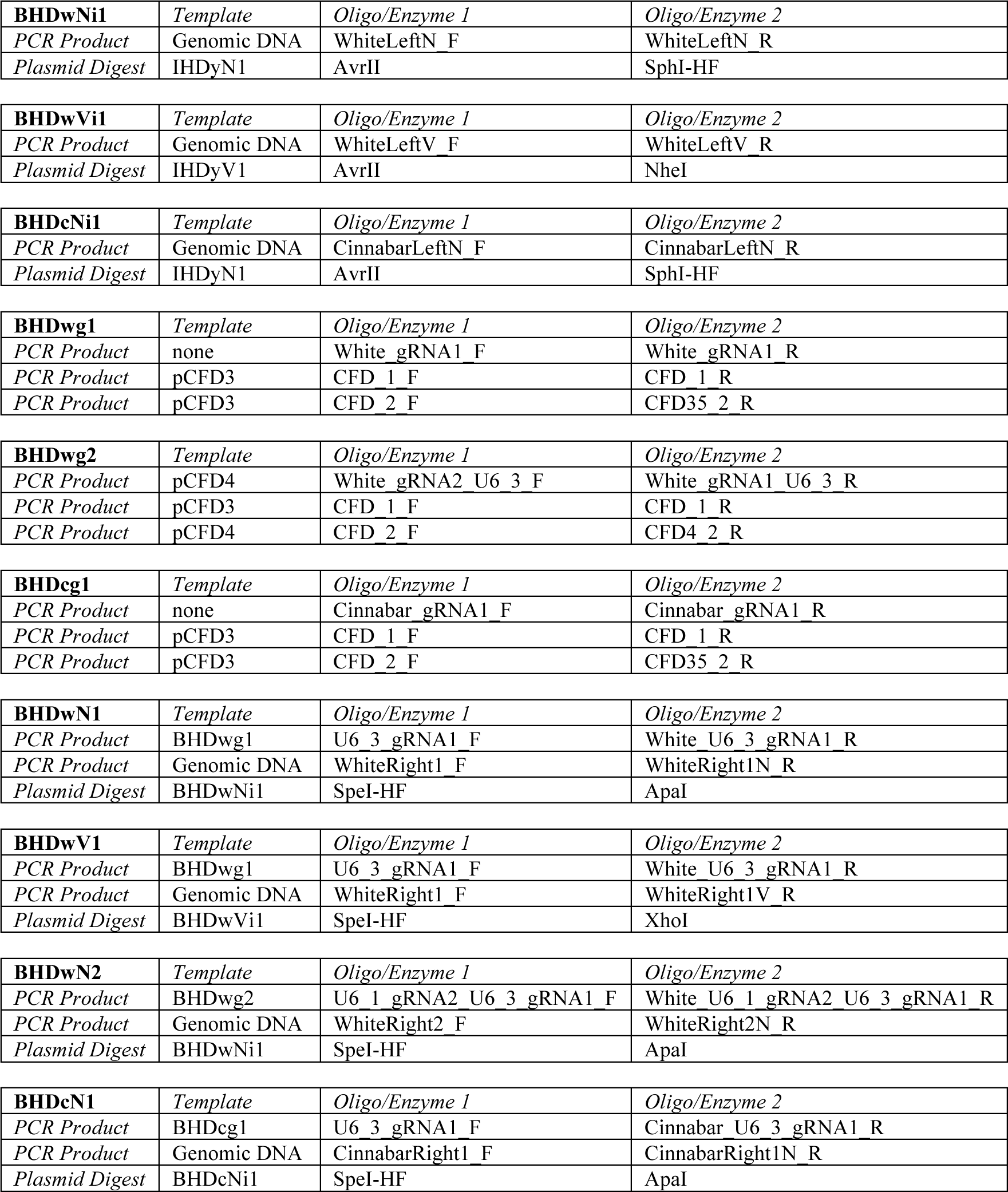

### Construction primer list

CFD_1_F: GTTTTAGAGCTAGAAATAGCAAGTTAAAATAAGG
CFD_1_R: GGCTATGCGTTGTTTGTTCTGC
CFD_2_F: AACAGTAGGCAGAACAAACAACGC
CFD35_2_R: CGACGTTAAATTGAAAATAGGTCTATATATACG
CFD4_2_R: CGAAGTTCACCCGGATATCTTTCCT
Cinnabar _gRNA1_F: TATATATAGACCTATTTTCAATTTAACGTCGCCACCGCCATACCCATGCG
Cinnabar _gRNA1_R: ATTTTAACTTGCTATTTCTAGCTCTAAAACCGCATGGGTATGGCGGTGGC
Cinnabar _U6_3_gRNA1_R: TGCCTCGC ACGCGTCCTGCAGGATGCATACGCATTAAGCGAACATT
CinnabarLeftN_F: AACCAATTCTGAACATTATCGCCTAGGGTACCACGGACAATCGCTTCAAATGGTTACACA
CinnabarLeftN_R: TTTCTCGAAAAGGGCCAGGAAGGAGCATGTCTAGAATGGGTATGGCGGTGGCCAGCA
CinnabarRight1_F: GTATGCATCCTGCAGGACGCGTGCGAGGCAGAATGCTCCACGATG
CinnabarRight1N_R:
GTTGTAAAACGACGGCCAGTCTTAAGCTCGGGCCGAGCTTTTGAGTAGTAGGTCCAGCCG
U6_1_gRNA2_U6_3_gRNA1_F:
GCTATACGAAGTTATAGAAGAGCACTAGTCGCGAATTTTCAACGTCCTCGATAGTATAGT
U6_3_gRNA1_F: ATGCTATACGAAGTTATAGAAGAGCACTAGGCTAGCTTTTTTGCTCACCTGTGATTGCTC
White _gRNA1_F: TATATATAGACCTATTTTCAATTTAACGTCGGCCAAAAGTTCGCCCGGAT
White _gRNA1_R: ATTTTAACTTGCTATTTCTAGCTCTAAAACATCCGGGCGAACTTTTGGCC
White _gRNA1_U6_3_R:
ATTTTAACTTGCTATTTCTAGCTCTAAAACATCCGGGCGAACTTTTGGCCGACGTTAAATTGAAAATAGGTCTAT
White_gRNA2_U6_3_F:
ATATATAGGAAAGATATCCGGGTGAACTTCGCATCCAAGTATCGCCATCCGTTTTAGAGCTAGAAATAGCAAG
White _U6_1_gRNA2_U6_3_gRNA1_R: ATCCCGGAACGCGTCCTGCAGGATGCATACGCATTAAGCGAACA
White _U6_3_gRNA1_R: TTCGCCCGACGCGTCCTGCAGGATGCATACGCATTAAGCGAACATT
White _U6_3_gRNA1_R: TTCGCCCGACGCGTCCTGCAGGATGCATACGCATTAAGCGAACATT
WhiteRight1_F: GTATGCATCCTGCAGGACGCGTCGGGCGAACTTTTGGCCGTGA
WhiteLeftN_F: ATTAACCAATTCTGAACATTATCGCCTAGGGTACCAGAGATTGAGTTTTCCCACCACCCA
WhiteLeftN_R: CGTTTCTCGAAAAGGGCCAGGAAGGAGCATGTCTAGAGATAGGCCACGCCGCAAACTGAG
WhiteLeftV_F: ATTAACCAATTCTGAACATTATCGCCTAGCCCGGGAGAGATTGAGTTTTCCCACCACCCA
WhiteLeftV_R: CCACCACACTGCTGCTCTTCGTGTTGGCTAGGTCGACGATAGGCCACGCCGCAAACTGAG
WhiteRight1_F: GTATGCATCCTGCAGGACGCGTCGGGCGAACTTTTGGCCGTGA
WhiteRight1N_R: TTGTAAAACGACGGCCAGTCTTAAGCTCGGGCCCTCCACTGGAACCACTCACCGTTGTC
WhiteRight1V_R: TCGCCCTTGAACTCGATTGACGGAAGAGCCTCGAGTCCACTGGAACCACTCACCGTTGTC
WhiteRight2_F: GTATGCATCCTGCAGGACGCGTTCCGGGATGCGACTGCTCAATG
WhiteRight2N_R: TTGTAAAACGACGGCCAGTCTTAAGCTCGGGCCCACTAAGAAGGGTGTGGAATCAGGCA

### Sequencing primer list

CinnabarLeft_S_F: TGCGAAAGCATAAATAGATTGTGGG
CinnabarLeft_S_R: TGAAGCTTAACTAGAATTATTGCCTGT
CinnabarRight_S_F: TGAGATCTTCGCTGGCATTCAG
CinnabarRight_S_R: ATGGACACCAGAAACTGTGGC
dsRed_S_F: CTGAAGGGCGAGATCCACAAG
gRNA1_S_F: TTGCTCACCTGTGATTGCTCC
IHD_S_R: TCTCGAAAATAATAAAGGGAAAATCAG
IHD_S_F: GGGTTATTGTCTCATGAGCGG
U6_1_gRNA2_S_F: CGAACATGGCCTTGGACGAAT
WhiteLeft_S_F: TGCACAGACGCCTTCATTTTT
WhiteLeft_S_R: TGCTCATCTAACCCCGAACAA
WhiteLeft_S2_F: CAGAGCTGCATTAACCAGGGCTTCG
WhiteRight_S_F: GAGAAAGGAAGCGTCTGGCAT
WhiteRight_S_R: TCGGAAGACGGCTGATGAATG
WhiteRight_S2_R: TCGGAAGACGGCTGATGAATGGTCA

**Data S1.**
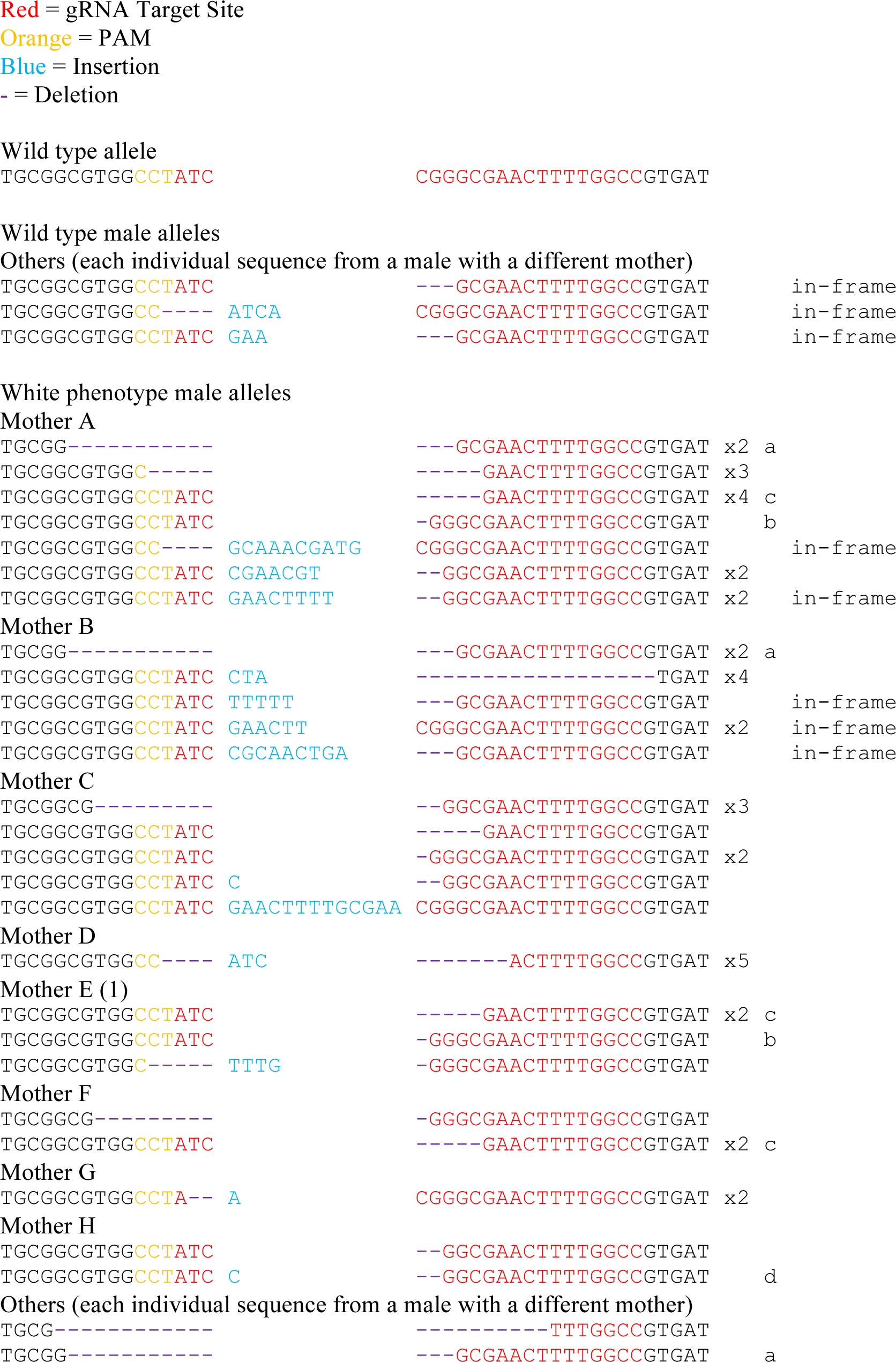
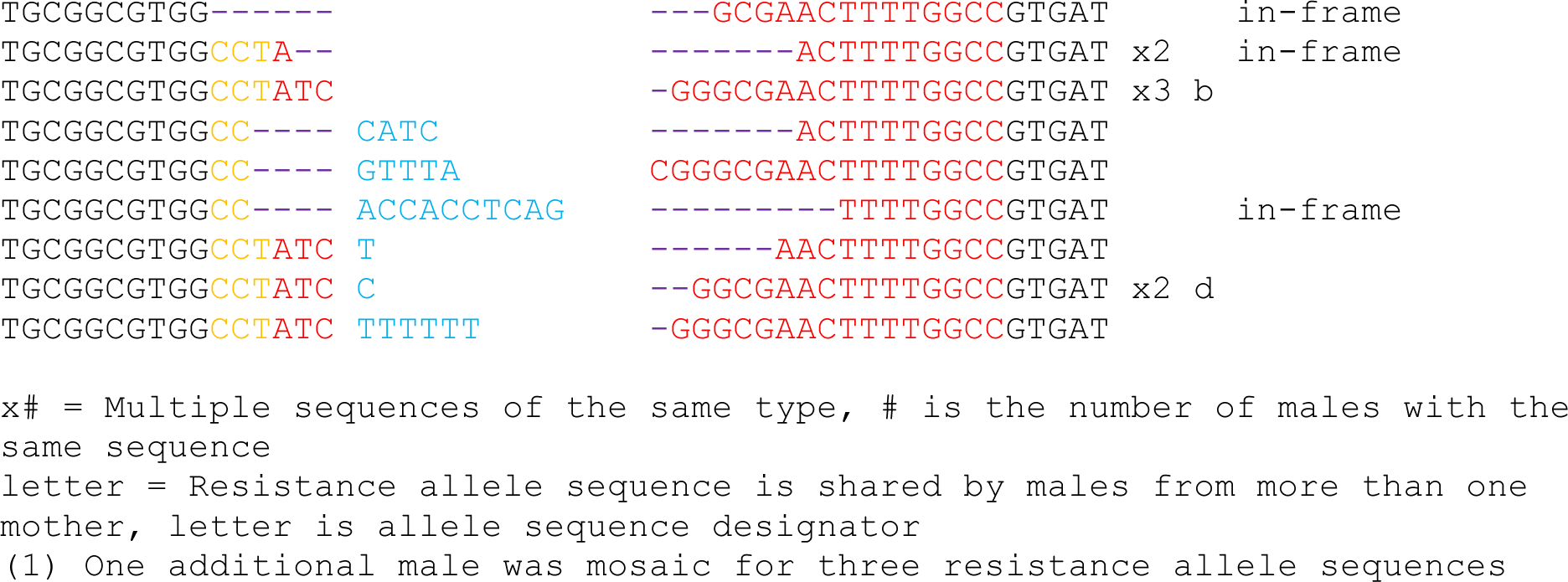
Resistance allele sequences from the *nanos* drive targeting *white*.

**Data S2.**
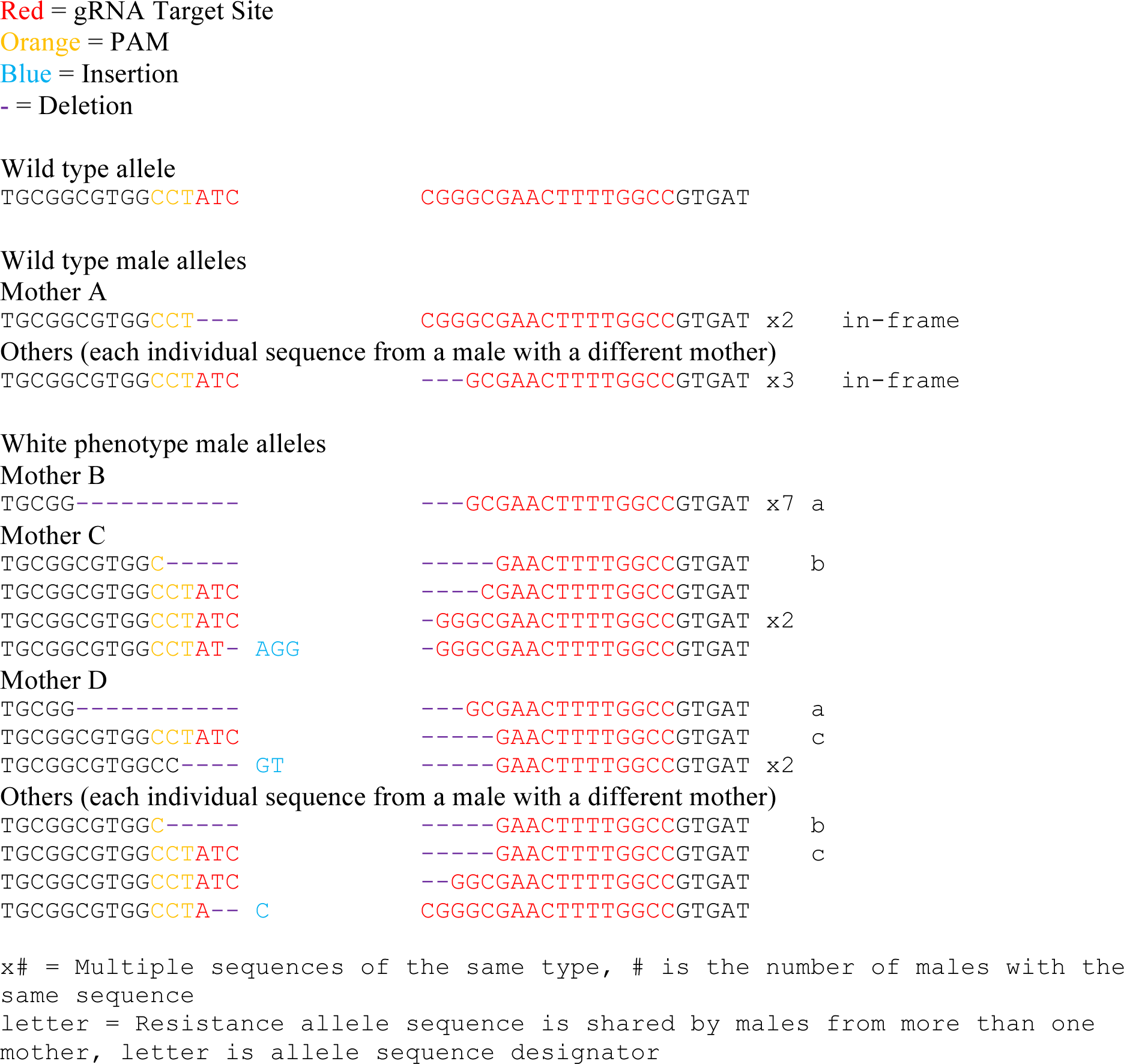
Resistance allele sequences from the *vasa* drive targeting *white*.

**Data S3.**
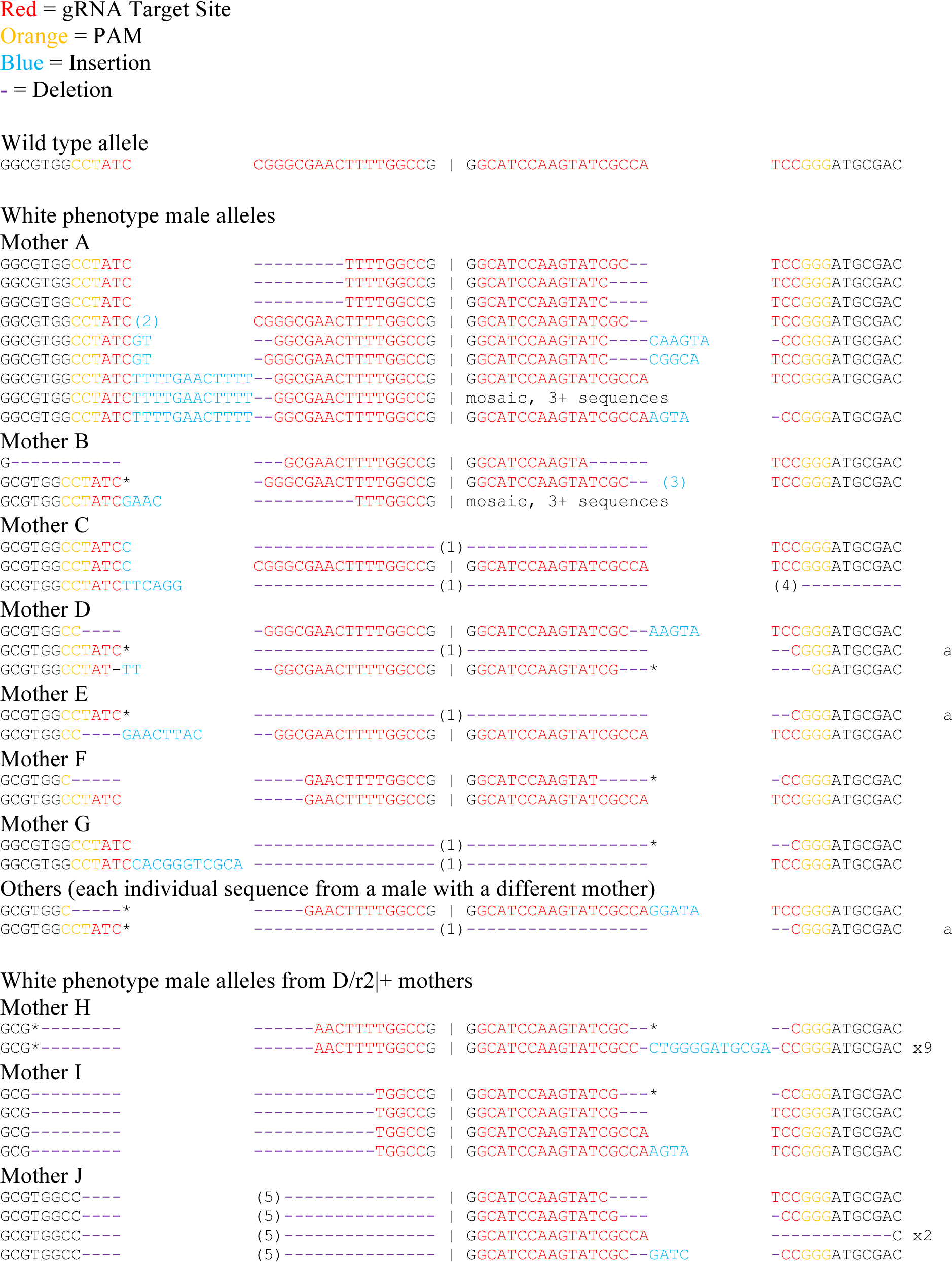
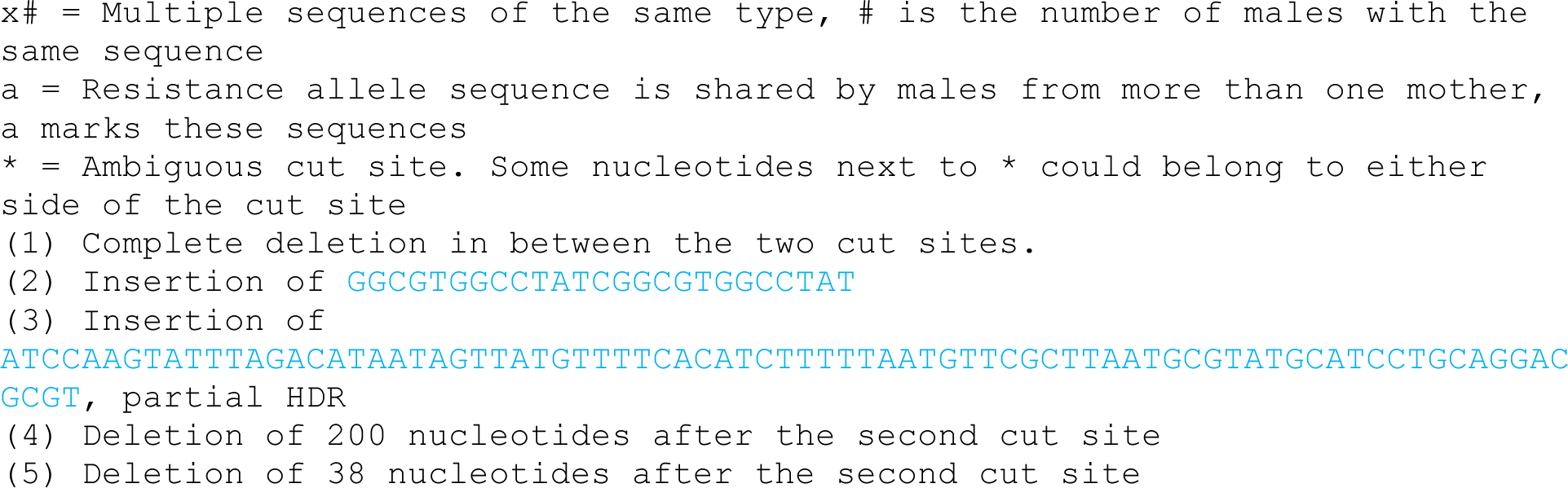
Resistance allele sequences from the two-gRNA *nanos* drive targeting *white*.

**Data S4.**
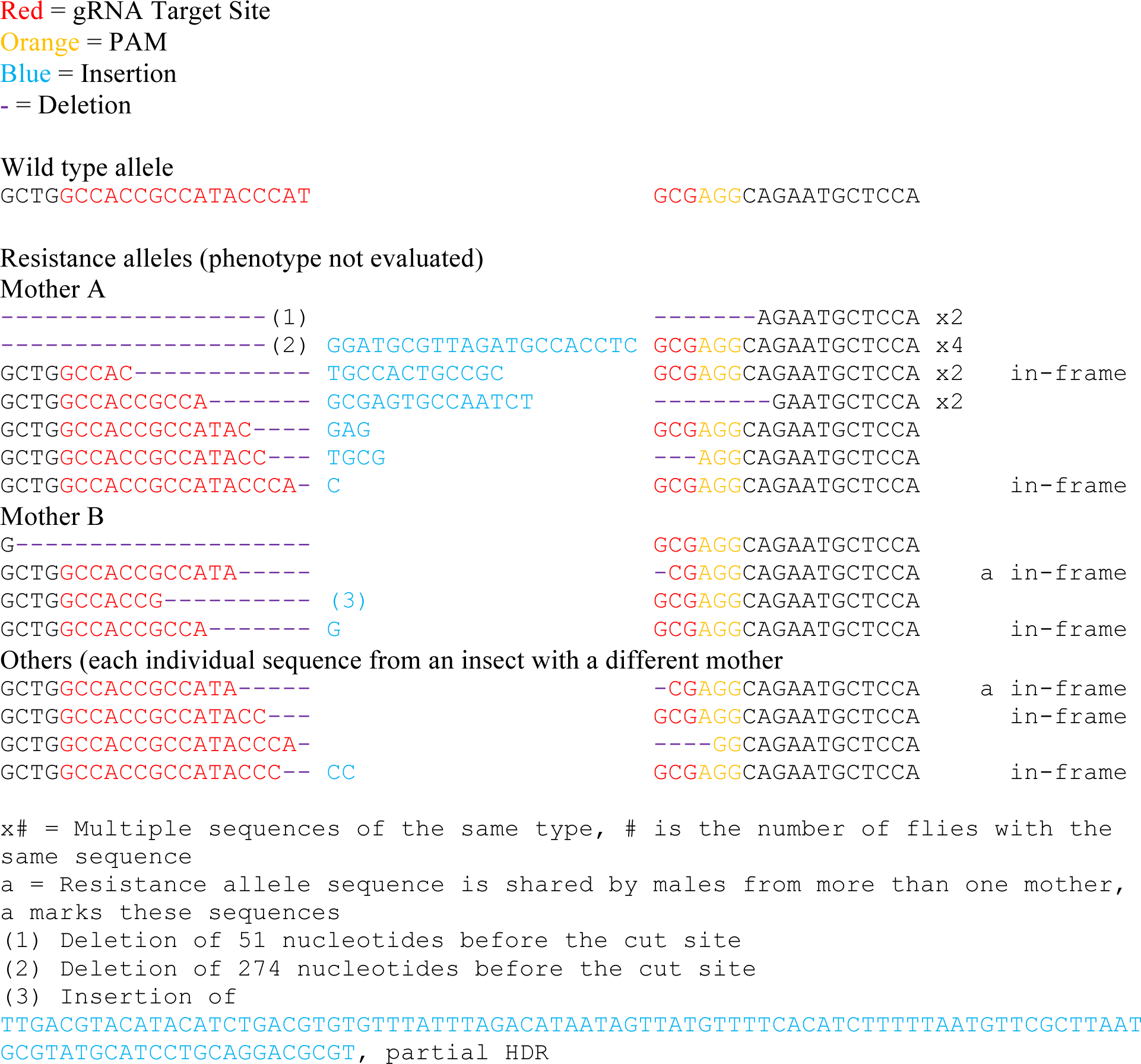
Resistance allele sequences from the *nanos* drive targeting *cinnabar*.

